# Remembering immunity: Neuronal ensembles in the insular cortex encode and retrieve specific immune responses

**DOI:** 10.1101/2020.12.03.409813

**Authors:** Tamar Koren, Maria Krot, Nadia T. Boshnak, Mariam Amer, Tamar Ben-Shaanan, Hilla Azulay-Debby, Haitham Hajjo, Eden Avishai, Maya Schiller, Hedva Haykin, Ben Korin, Dorit Cohen-Farfara, Fahed Hakim, Kobi Rosenblum, Asya Rolls

**Affiliations:** Department of Immunology, Rappaport Faculty of Medicine, Technion - Israel Institute of Technology, Haifa, Israel; Department of Neuroscience, Rappaport Faculty of Medicine, Technion - Israel Institute of Technology, Haifa, Israel; Department of Microbiology and Immunology, University of California, San Francisco, San Francisco, CA, USA; Howard Hughes Medical Institute, University of California, San Francisco, San Francisco, CA, USA; Pediatric Pulmonary Unit, Rambam Health Care Campus, Haifa, Israel; Cancer Research Center, EMMS Hospital, Nazareth, Israel; Sagol Department of Neurobiology, University of Haifa, Haifa, Israel; Center for Gene Manipulation in the Brain, University of Haifa, Haifa, Israel

## Abstract

Increasing evidence indicates that the brain regulates peripheral immunity. Yet, it remains unclear whether and how the brain represents the state of the immune system. Here, we show that immune-related information is stored in the brain’s insular cortex (InsCtx). Using activity-dependent cell labeling in mice (*Fos*^*TRAP*^), we captured neuronal ensembles in the InsCtx that were active under two different inflammatory conditions (DSS-induced colitis and Zymosan-induced peritonitis). Chemogenetic reactivation of these neuronal ensembles was sufficient to broadly retrieve the inflammatory state under which these neurons were captured. Thus, we show that the brain can encode and initiate specific immune responses, extending the classical concept of immunological memory to neuronal representations of immunity.

## Main Text

Accumulating data indicate that the brain can affect immunity, as evidenced for example, by the effects of stress ^1–3^, stroke ^4–6^ and reward system activity ^7,8^ on the peripheral immune system. However, our understanding of this neuro-immune interaction is still limited. Importantly, we do not know how the brain evaluates and represents the state of the immune system.

As a central regulator, it is likely that the brain receives feedback from any system it controls, and by extension, forms representations of the immune system. This concept is supported by several lines of evidence: First, anatomically, the brain receives immune-related information via sensory inputs from peripheral immune organs, such as the spleen and lymph nodes ^9–11^. Second, the brain responds to peripheral immune challenges. For example, neuro-imaging studies identified increased activation of specific brain regions (e.g., the amygdala, hypothalamus, brainstem, thalamus and insular cortex) during peripheral inflammation ^12,13^. Finally, the immune system has been shown to respond to behavioral conditioning by associating a specific immune response (e.g., immune suppression) with an external cue (e.g., saccharine water), such that presentation of the cue alone can elicit the immune response ^14,15^. This suggested that the brain must encode immune-related information ^16^. Furthermore, lesion studies identified specific brain regions required for immune conditioning, notably, the insular cortex (InsCtx) ^17,18^. However, the existence of such immune representations in the brain was never directly demonstrated.

The current study was designed to test the existence of immune-related representations in the brain and to determine their relevance to immune regulation. We hypothesized that the InsCtx, and specifically the posterior InsCtx, is especially suited to contain such a representation of the immune system. This region is considered the primary cortical site of interoception, i.e., the sensing of the body’s physiological state ^19,20^, and integrates information regarding bodily sensations, such as pain, hunger and visceral signals ^21^. Although immune-related information is not a conventional aspect of interoception, it could provide an important indication of the organism’s physiological status ^22^, and thus, may also be processed in the posterior InsCtx. Moreover, the InsCtx is well positioned to gather immune-related information, as it receives input from peripheral neurons that respond to immune signals ^9,23,24^. Accordingly, studies have shown that immune challenges impact insular activity ^25,26^. Finally, as previously mentioned, lesion studies have shown that the InsCtx is essential for immune conditioning ^27,28^, further supporting the hypothesis that this brain region can form immune-related representations.

To test this hypothesis, we used transgenic targeted-recombination-in-active-populations (TRAP) mice 29,30 to capture neurons that were active during a peripheral immune challenge (**Figure 1A**). These mice express iCreER^T2^ under the control of an activity-dependent c-Fos promoter (*Fos*^*TRAP*^ mice), which serves as an indicator of neuronal activity. In the presence of Tamoxifen, active neurons drive Cre-dependent recombination to induce the expression of an effector gene (e.g., fluorescent reporter). For the immune challenge, we chose a model of colon inflammation (colitis). Emerging experimental and epidemiological data established the significance of the gut-brain axis in gastrointestinal (GI) inflammatory conditions ^31–35^. The InsCtx receives information through the brainstem and thalamus from this highly innervated and immunologically active visceral organ ^36,37^. Thus, we used a model of inflammatory bowel disease (IBD), DSS-induced colitis, to capture its potential representation in the InsCtx.

**Figure 1.**
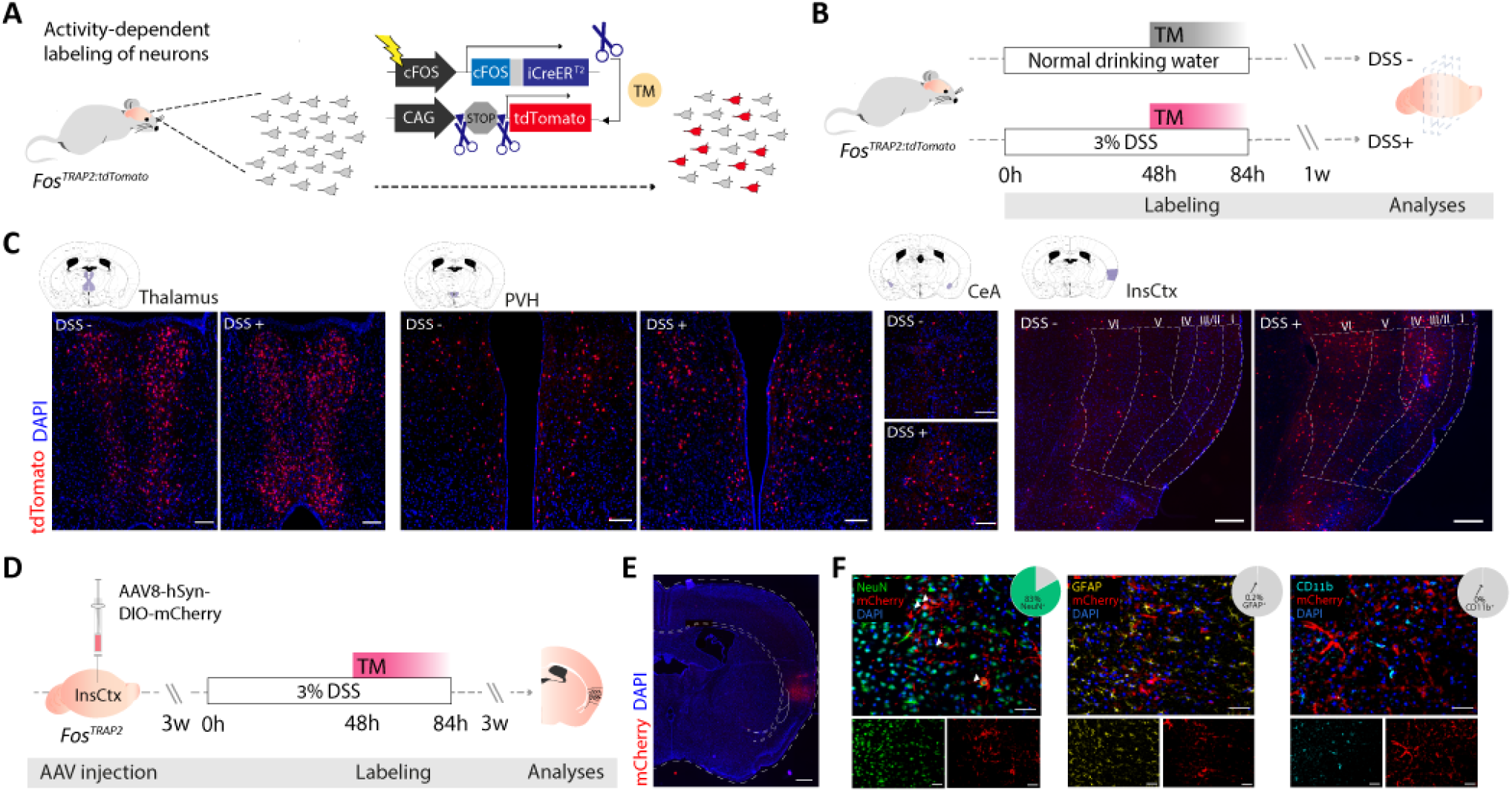
Activity-dependent labeling of neurons during DSS-induced colitis in the mouse InsCtx. (**A**) Schematic representation of activity-dependent neuronal labeling in *Fos*^*TRAP2:tdTomato*^ mice. iCreER^T2^ is expressed in activated neurons, indicated here by cFos expression. The administration of Tamoxifen (TM) allows iCreER^T2^ recombination to occur only in active neurons. Thus, only neurons that are active while TM is present at sufficient levels (~24-36 hours following administration) will express the Cre-dependent fluorescent reporter (tdTomato). **(B)** Experimental design for fluorescently labeling neurons active during DSS-induced colitis (DSS+) or a normal drinking water regime (DSS-) in *Fos^TRAP2:tdTomato^* mice. **(C)** Fluorescence microscopy imaging showing the expression of tdTomato in different brain regions of both experimental groups: Thalamus, Paraventricular nucleus of the hypothalamus (PVH), Central amygdala (CeA) and right Insular cortex (InsCtx), from left to right (*n*=3, 5). Scale bars: 50 μm (Thalamus, PVH, CeA), 100 μm (InsCtx). **(D)** Experimental design for fluorescently labeling neurons active during DSS-induced colitis in *Fos^TRAP2^* mice. **(E)** Fluorescence microscopy imaging showing the expression of the injected Cre-dependent fluorescent marker (mCherry, in red) in the InsCtx, 3 weeks following TM administration. Scale bar: 250 μm. **(F)** Representative brain micrographs showing immunofluorescence (IF) staining of the labeled cells in the InsCtx (mCherry, red) co-stained for neuronal (NeuN, green; *n*= 6), astrocytic (GFAP, yellow; *n*= 5) and microglial (CD11b, cyan; *n*= 6) markers. Arrows indicate double-positive cells for mCherry and NeuN. Pie charts indicate percentages of double stained neurons out of all mCherry^+^ cells. Scale bars: 50 μm. AAV, adeno-associated virus; lightning symbol, neuronal activity; TM, Tamoxifen; DSS, dextran sulfate sodium; InsCtx, insular cortex. Data represent two independent repeats.

We crossed the TRAP mice with a Cre-dependent tdTomato reporter line, *Ai14D* ^38^ to visualize the captured cells. Mice were injected with Tamoxifen (inducing Cre recombination and tdTomato expression) 48h after initiation of the DSS treatment (DSS+ group). As controls, we used another group of TRAP-tdTomato mice that was injected with Tamoxifen at the same time, but did not receive DSS (DSS-group; **Figure 1B**). This group allowed us to compare between inflammation-related activity in the brain, and its activity during healthy homeostasis. Fluorescence microscopy of brain cryosections revealed increased tdTomato expression (indicative of neuronal activity) in several brain areas that were expected to respond to aberrant physiological states such as colon inflammation. Notable regions included the thalamus, which receives sensory and visceral information, the paraventricular hypothalamic (PVH) nuclei, known to play a major role in maintaining the physiological homeostasis and stress responses, the central amygdala (CeA), which is associated with fear and memory formation, and the InsCtx (**Figure 1C**). These findings are consistent with studies indicating that these brain areas, including the InsCtx, react to peripheral immune processes ^13,39–42^.

Based on this correlative evidence and our prediction of the InsCtx functioning as an interoceptive site that receives immune-related information, we chose to focus on this area. Moreover, since the InsCtx exhibits some functional lateralization, we thus focused on the right InsCtx, which is more dominantly associated with sympathetic afferent innervation and visceral information processing ^43^. To target the TRAPing to this brain region and limit the labeling to neuronal populations, we used a viral vector instead of the TRAP-tdTomato reporter. The viral vector encoded the mCherry fluorescent reporter, expressed in a Cre-dependent manner under a neuronal-specific promoter, Synapsin, and was stereotactically injected to the right posterior InsCtx (**Figure 1D**). To capture active neurons during colitis, we injected mice with Tamoxifen (inducing Cre recombination and mCherry expression in c-Fos-expressing neurons) 48h after initiation of the DSS treatment (**Figure 1D**). Immunofluorescence (IF) staining revealed that, as expected (based on the use of Synapsin promoter), most of the labeled cells (**Figure 1E**) were indeed neurons, rather than astrocytes or microglia (**Figure 1F**). Thus, based on the *Fos*^*TRAP:tdTomato*^ data (**Figure 1C**) and these virus-based labeling experiments, we concluded that DSS-induced colitis is associated with increased activation of neurons in the InsCtx. However, these correlative findings are not sufficient to demonstrate that these neurons indeed encode immune-relevant information.

To directly address this question, we sought to reactivate the TRAPed neurons and evaluate their effects on the immune response. To this end, we used *Fos*^*TRAP*^ mice to capture the activated neurons. For neuronal reactivation, we co-expressed a stimulatory DREADD (designer receptors exclusively activated by designer drugs; Gq) in the captured neurons, in addition to the fluorescent reporter. DREADDs are mutated muscarinic G protein-coupled receptors (GPCRs) activated by a synthetic ligand, clozapine *N*-oxide (CNO). Thus, CNO application enables the reactivation of the captured neurons at will. We chose DREADDs rather than optogenetics to control neuronal activity due to their more extended time frame of activity, which is more suitable for the time scale of immune reactions. The control group was injected with a virus encoding the mCherry fluorescent reporter, but lacking the DREADD-coding sequence (Sham control, **Figure 2A**). This sham group allowed us to control for the surgical procedure, viral vector expression, and CNO application. As in the previous experiment, we injected Tamoxifen 48h following initiation of the DSS treatment to capture the active neurons and enable their expression of both Gq-DREADDs and a fluorescent reporter. The mice were allowed to recover from the DSS-induced colitis for 4 weeks (based on previous studies indicating of inflammation resolution at this time point ^44^) and were then injected with CNO to re-activate the captured neuronal ensembles; the effects of this activation on the immune response were then evaluated.

**Figure 2.**
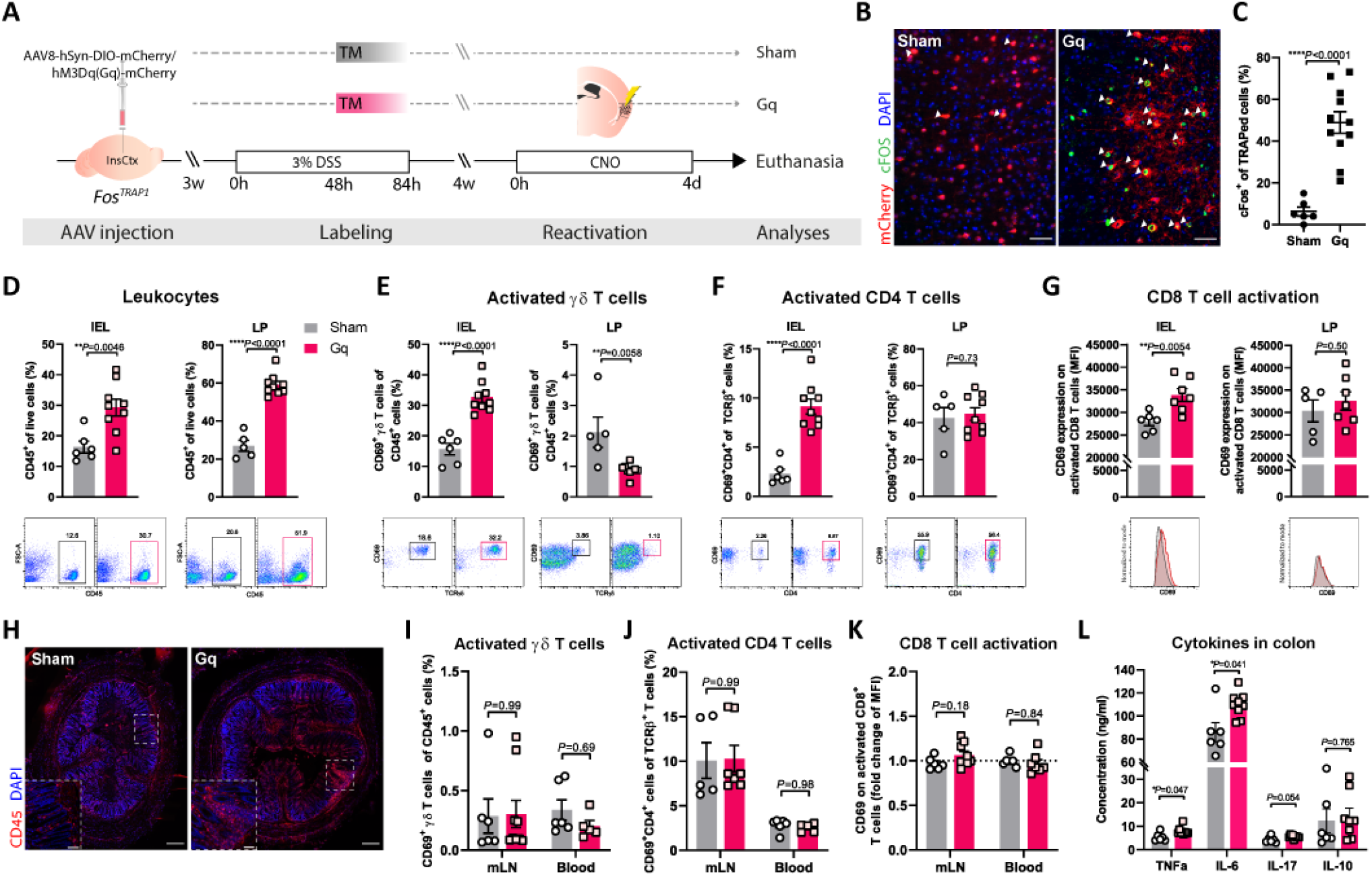
Reactivation of InsCtx neuronal ensembles captured during colitis, recapitulates an inflammatory state in the colon. (**A)** Schematic representation of the protocol for capturing neuronal ensembles that were active during DSS-induced colitis in the *FosTRAP1* mice, and reactivation of the captured neurons after recovery from the inflammation. Mice were injected with the viral vector expressing the coding sequence for the active form of DREADD (Gq) or a sham virus (carrying only the fluorescent reporter, mCherry). Mice were treated with DSS 3 weeks after surgery then injected with TM (48 h after the beginning of DSS administration). Upon cessation of DSS treatment, mice were allowed to recover from the inflammation (4 weeks) and were then injected with CNO for 4 days (at 24 h intervals) to reactivate the captured neuronal ensembles. Controls were also treated with CNO, but since they did not express DREADDs, the captured neurons were not activated in these mice. Finally, mice were euthanized, and their colons, mesenteric lymph nodes and peripheral blood were collected for immunological analyses. **(B)** Representative micrographs of InsCtx showing both sham viral vector-(left, mCherry) and Gq-DREADD-(right, mCherry) expressing neurons, immunofluorescently (IF) stained for c-Fos (green) following CNO administration. Scale bar: 100 μm. **(C)** Quantification of c-Fos-expressing neurons (IF-stained) percentages out of mCherry-expressing neurons in the InsCtx. Data are represented as mean±s.e.m from *n*=5, 11 mice; Student’s *t*-test. **(D-G)** Graphs derived from flow cytometry analysis showing percentages of **(D)** leukocytes (CD45^+^ cells; *n*=6/5,9), **(E)** activated γδ T cells (*n*=6/5,9), **(F)** activated CD4^+^ T cells (*n*=6/5,9), and **(G)** CD69 expression (MFI) on CD8^+^ T cells (*n*=6/5,6/7) in the colonic mucosal layer (IEL, LP) of Gq mice relative to their sham viral vector controls following neuronal reactivation. Data from individual mice are shown and values are represented as mean±s.e.m; Student’s *t*-test. **(H**)Representative micrographs of colon tissue (*n*=6, 8), IF-stained for CD45 (red) and DAPI (blue). Scale bars: 100 μm (whole section), 50 μm (partial section). **(I-J)** Graphs showing percentages of mesenteric lymph nodes (mLN; *n*=5/6, 9/7) and peripheral blood (*n*=6, 9) **(I)** activated γδ T cells, **(J)** activated CD4^+^ T cells, and **(K)** CD69 expression (MFI) on CD8^+^ T cells, the same immune cell parameters that significantly changed in the colon of the Gq group compared to the sham group. Data from individual mice are shown and values are represented as mean±s.e.m; Multiple *t*-tests. **(L)** Graph derived from cytokine ELISA showing concentrations of TNFα, IL-6, IL-17, and IL-10 in colons of the Gq group compared to its sham viral vector controls. Data are represented as mean±s.e.m from *n=* 6/5, 9 mice; Multiple *t*-tests. One sample was excluded from the data; ROUT method for outlier detection (Q=1%). ELISA, enzyme-linked immunosorbent assay; InsCtx, insular cortex; TM, Tamoxifen; DSS, dextran sulfate sodium; lightning symbol, neuronal activity; CNO, clozapine *N*-oxide; IEL, intraepithelial lymphocytes; LP, lamina propria; MFI, median fluorescence intensity. Data represent two pooled independent repeats.

Since the time elapsing between the neuronal TRAPing and reactivation could potentially affect the efficiency of the neuronal reactivation, we validated that our DREADD manipulation maintained the ability to induce neuronal activation. Immunofluorescence staining for the activation marker, c-Fos (**Figure 2B**), indicated that, indeed, significantly more neurons (*P*<0.0001) were active in the Gq group (48.9% ± 5.1 s.e.m c-Fos^+^ cells) compared to sham controls (6.33% ± 2.17 s.e.m) following CNO application (**Figure 2C**). To analyze the functional outcomes of this neuronal reactivation on the inflammatory response in the colon, we evaluated several immune parameters in the colon tissue. We found that even in the absence of an additional immune challenge, neuronal reactivation alone resulted in heightened immune activity in the colon of the Gq-mice compared to their sham controls. Specifically, there was a significant increase of the percentage of leukocytes (CD45^+^ cells) present in the mucosal layers of the colon, including the intraepithelial-lymphocytes (IEL; *P*=0.0046) and leukocytes within the lamina propria (LP; *P*<0.0001; **Figure 2D, Figure S1**). Percentages of activated γδ T cells increased in the IEL (*P*<0.0001), and decreased in the LP (*P*=0.0058), in the Gq group compared to the sham treated mice (**Figure 2E, Figure S1**), a characteristic trend that was previously <u>demonstrated for</u> γδ T cells during intestinal infection ^45^. Moreover, in the IEL, we found higher proportions of activated CD4^+^ T cells (CD69^+^CD4^+^, *P*<0.0001; **Figure 2F, Figure S1**) and an increase in the activation of CD8^+^ T cells (*P*=0.0043; **Figure 2G, Figure S1**). The increase in leukocyte presence in the colonic mucosal layer, observed via flow cytometry, was further supported by the IF analysis of the colon tissue in both groups (*n*=6, 8, **Figure 1H**). Importantly, these brain-induced effects were limited to the GI tract, as we did not detect any significant changes in immune-cell populations in the peripheral blood or the mesenteric lymph nodes (mLN) (**Figures 2I-K**).

Cytokine levels in the colon were also elevated in the Gq-group, compared to their sham viral vector controls. Specifically, the pro-inflammatory cytokines, TNFα and IL-6, were significantly increased in the colon (*P*=0.047, *P*=0.041, respectively). There were no significant changes in the levels of IL-17 (*P*=0.054) or IL-10 (*P*=0.765; **Figure 2L**). Collectively, these data suggest that reactivation of the InsCtx neurons that were active during the original inflammation was sufficient to induce an inflammatory response, specifically in the colon.

Nevertheless, as controls, we used mice that only express the fluorescent reporter, and thus, CNO application did not induce any neuronal activation in this group. Therefore, it may be argued that this experiment reflects the impact of general InsCtx activation on immune activity. To directly test whether specific neuronal ensembles in the InsCtx can differentially affect the immune response, specifically in the colon, we performed an additional experiment in which all mice were injected with a Gq-DREADD. Accordingly, we were able to induce neuronal reactivation in all experimental groups. However, by varying the timing of Tamoxifen administration, we could capture neuronal activity at different time points relative to colitis induction. In this experiment, one group of mice received Tamoxifen injection before DSS (Gq-before), a second group received Tamoxifen injection during the inflammatory response (Gq-during), and the third group was injected with Tamoxifen during the recovery stage (Gq-after). Thus, all three groups were subjected to the same experimental procedures (injected with the same viral vector, and treated with DSS). After the mice recovered from the initial DSS-induced inflammation, we injected all three groups of mice with CNO to reactivate the captured neuronal populations, and evaluated the presence of colon inflammation (**Figure 3A**).

**Figure 3.**
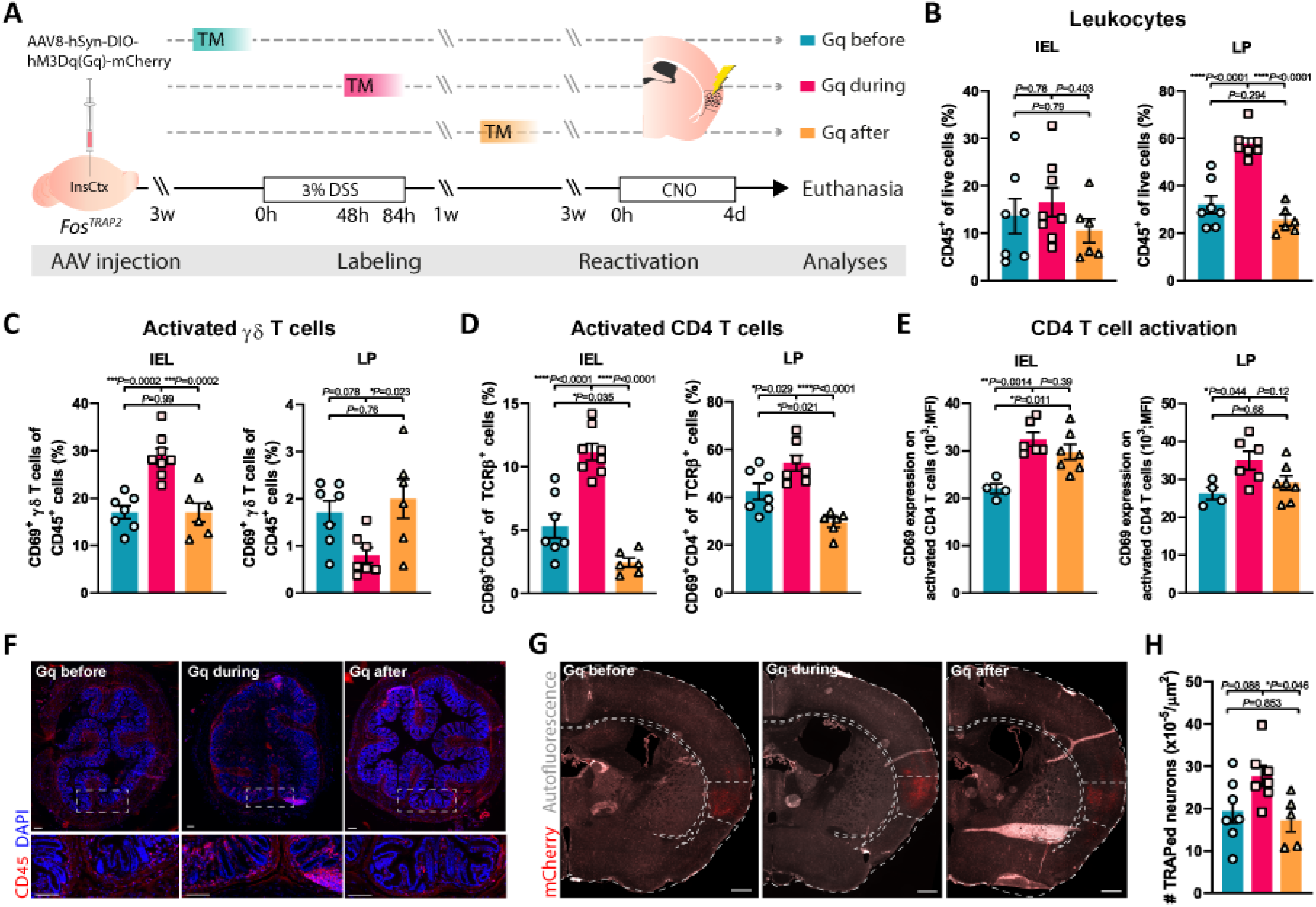
Reactivation of neuronal ensembles captured at different stages of DSS induced colitis generate different immune responses in the colon. (**A**) Schematic representation of enabling DREADD expression in neuronal ensembles that were active before, during or after DSS-induced colitis in the InsCtx of *Fos^TRAP2^* mice (via different timing of TM injections, relative to DSS treatment). Four weeks after cessation of DSS treatment, all mice received daily CNO injections for 4 days, and the immune response was analyzed. (**B-E**) Graphs derived from flow cytometry analysis showing percentages of (**B**) leukocytes (CD45^+^ cells), (**C**) activated ɣɗ T cells (**D**) activated CD4^+^ T cells, and (**E**) CD69 expression (MFI) on CD4^+^ T cells in the mucosal layer of the colon (IEL, LP) in all three Gq groups, labeled before (Gq before), during (Gq during), and after (Gq after) inflammation. Data from individual mice are shown, and values are represented as mean±s.e.m from *n*=7, 8/7, 6 (**B-D**), *n*=4, 6, 7 (**E**) mice; One-way ANOVA and Multiple *t*-tests. (**F**) Representative micrographs of colon tissue (*n*=6/5,9), IF-stained for CD45 (red) and DAPI (blue). Scale bars: 50 μm. (**G**) Representative fluorescence microscopy imaging showing the expression of the injected Cre-dependent DREADD (mCherry, in red) in the InsCtx of all 3 experimental groups. Scale bar: 250 μm. (**H**) Quantification of DREADD-expressing neurons in the InsCtx of all 3 experimental groups. Data are represented as mean±s.e.m from *n*= 7, 7, 5 mice; One-way Anova and Multiple *t*-tests. InsCtx, Insular cortex; TM, Tamoxifen; DSS, dextran sulfate sodium; CNO, clozapine-*N*-oxide; IEL, intraepithelial lymphocytes; LP, lamina propria; MFI, median fluorescence intensity. Data represent two pooled independent repeats.

As mentioned above, all groups expressed the Gq-DREADD. However, following CNO application, only the Gq-during group (labeled during the inflammation), showed a significant increase in the inflammatory response in the colon. As in the previous experiment (**Figure 2D-H**), in the Gq-during group, we detected an overall increase in the leukocyte population (CD45^+^ cells) within the LP (*P*<0.0001; **Figure 3B, Figure S2A-B**). Percentages of activated γδ T cells in the IEL population increased (*P*<0.0001), while activated γδ T cells in the LP decreased (*P=*0.0083; **Figure 3C, Figure S2C-D**). Activated CD4^+^ T cells (CD69^+^CD4^+^) increased in the IEL and LP (*P*<0.0001, *P*=0.0081, respectively; **Figure 3D, Figure S2E-F**), as well as their extent of activation (*P*=0.0017, *P*=0.0395, respectively; **Figure 3E, Figure S2G-H**). IF analysis of the colonic tissue, revealed similar effects (*n*=3, 4, 3, **Figure 3F**). Hence, an elevated immune response was evident only upon the reactivation of neurons captured during the original inflammation.

Although the neuronal reactivation induced an inflammatory response, some of the observed effects were different (in magnitude or direction) from the response directly induced by the initial DSS administration (as observed at the time of TRAPing; **Figure S3A-B**). For example, IFNγ-expressing leukocytes and CD103^-^CD11b^+^ dendritic cells increased in the colon in both the initial response and the reactivation. Yet, other effects, such as those found on γδ T cells and total percentage of leukocytes, were induced by InsCtx reactivation alone (**Figure S3C**). The similarities between the original immune response and the response induced by neuronal reactivation, suggest that immune-related information is encoded in the brain. However, the differences between these responses, highlight the potential limitations of our experimental paradigm, the complexity of such brain representations and the interaction with the peripheral tissue.

Our findings indicate a causal relationship between the activation of specific neuronal ensembles captured during DSS-induced colitis and the induction of an inflammatory response in the colon. Yet, as shown in Figure 3G-H, more neurons were captured in the Gq-during group compared to the Gq-before and Gq-after groups. Thus, it is possible that the difference in the immunological outcome between these experimental groups is a manifestation of the number of activated neurons rather than the specific information they encode. Namely, the InsCtx may innately induce an immune response, but only when a sufficient number of neurons are activated, does this effect become evident. To control for this possibility and target more neurons in the InsCtx, we expressed DREADDs in a Cre-independent manner, which allowed us to target approximately twice (2.06±0.42 fold; *P*=0.0045) the number of neurons in the same region in the InsCtx (**Figure 4A-E**), irrespective of prior activation. Nevertheless, this robust yet general activation, did not induce any inflammatory effects in the colon (**Figure 4F-J**), suggesting that unique immune-related information is carried by specific neuronal ensembles in the InsCtx.

**Figure 4.**
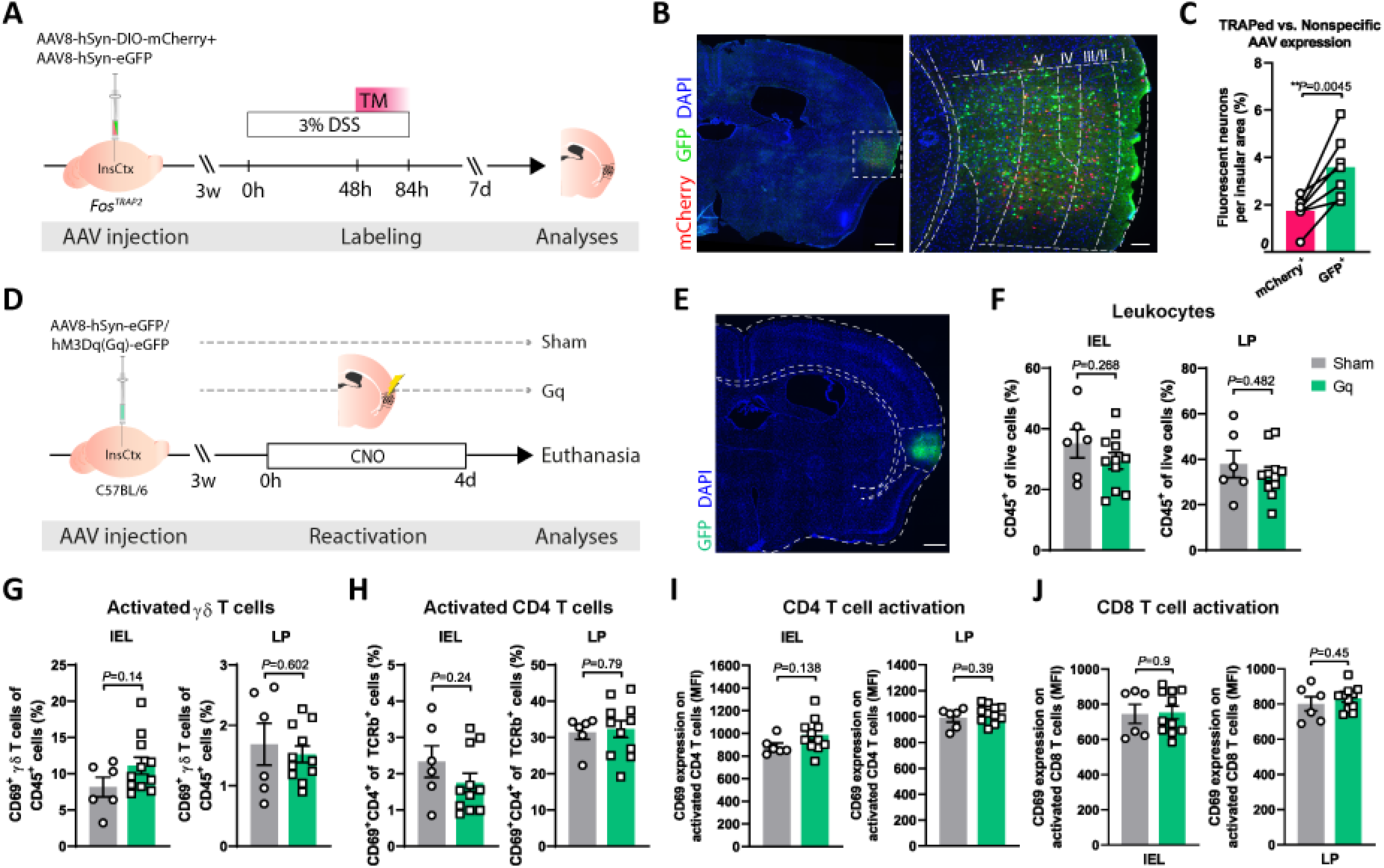
Nonspecific activation of the InsCtx elicits no apparent cellular immune response in the colon. (**A**) Experimental design for fluorescently labeling neurons in both a Cre-dependent (mCherry) and Cre-independent (GFP) manner in *Fos^TRAP2^* mice (**B**) Representative fluorescence microscopy imaging showing the expression of the injected fluorescent markers in the InsCtx, 7 days following TM administration. Scale bars: 250 μm (left), 100 μm (right). (**C**) Quantification of activity-dependent and nonspecific labeled neurons (mCherry and GFP, respectively) in the InsCtx. Data are represented as mean±s.e.m from *n*= 7 mice; Paired *t*-test. (**D**) Schematic representation of viral vector expression in InsCtx neurons in a nonspecific manner (Cre-independent) in C57BL/6 mice. Three weeks after stereotactic injections, all mice received daily CNO injections for 4 days, and the immune response was analyzed. (**E**) Representative fluorescence microscopy imaging showing the expression of the injected DREADD (GFP, in green) in the InsCtx. Scale bar: 250 μm. (**F-I**) Graphs derived from flow cytometry analysis showing percentages of (**F**) leukocytes (CD45^+^ cells), (**G**) activated ɣɗ T cells (**H**) activated CD4^+^ T cells, and CD69 expression (MFI) on (**I**) CD4^+^ and (**J**) CD8^+^ T cells in the mucosal layer of the colon (IEL, LP) of Gq mice relative to their sham viral vector controls following neuronal reactivation. Data from individual mice are shown, and values are represented as mean±s.e.m from *n*=6, 11 mice; Student’s *t*-test. Insular cortex; TM, Tamoxifen; CNO, clozapine-*N*-oxide; IEL, intraepithelial lymphocytes; LP, lamina propria; MFI, median fluorescence intensity. Data represent two pooled independent repeats.

The observed neuronal reactivation effects on immunity may be specific to the gut-brain axis, as so far, we analyzed a single type of immune response, colitis, in our experiments. To directly examine the degree of specificity with which the immune information can be encoded by neuronal ensembles in the InsCtx, we tested an unrelated immune challenge. We chose the Zymosan-induced peritonitis (ZIP), a well-defined inflammatory process affecting a distinct visceral area (the peritoneal cavity). Although it is an inflammatory process as well, this model is anatomically and immunologically distinct from the DSS-induced inflammatory model. As in the previous experiments, *Fos*^*TRAP*^ mice were stereotactically injected with the Gq-DREADD or a sham viral vector to their InsCtx. One Gq group received Tamoxifen before ZIP (Gq-before), and a second Gq group (Gq-during) and the sham viral vector-expressing group (Sham) were injected with Tamoxifen during ZIP (**Figure 5A**). In contrast to the DSS-induced colitis, there was no difference in the number of the TRAPed neurons captured during ZIP compared to their controls (*P*=0.723, **Figure 5B-C**), possibly because of the more acute nature of this model. Following recovery, all groups were injected with a single dose of CNO to induce neuronal reactivation in the Gq-groups, corresponding to the single injection of Zymosan applied during TRAPing (**Figure 5A**).

**Figure 5.**
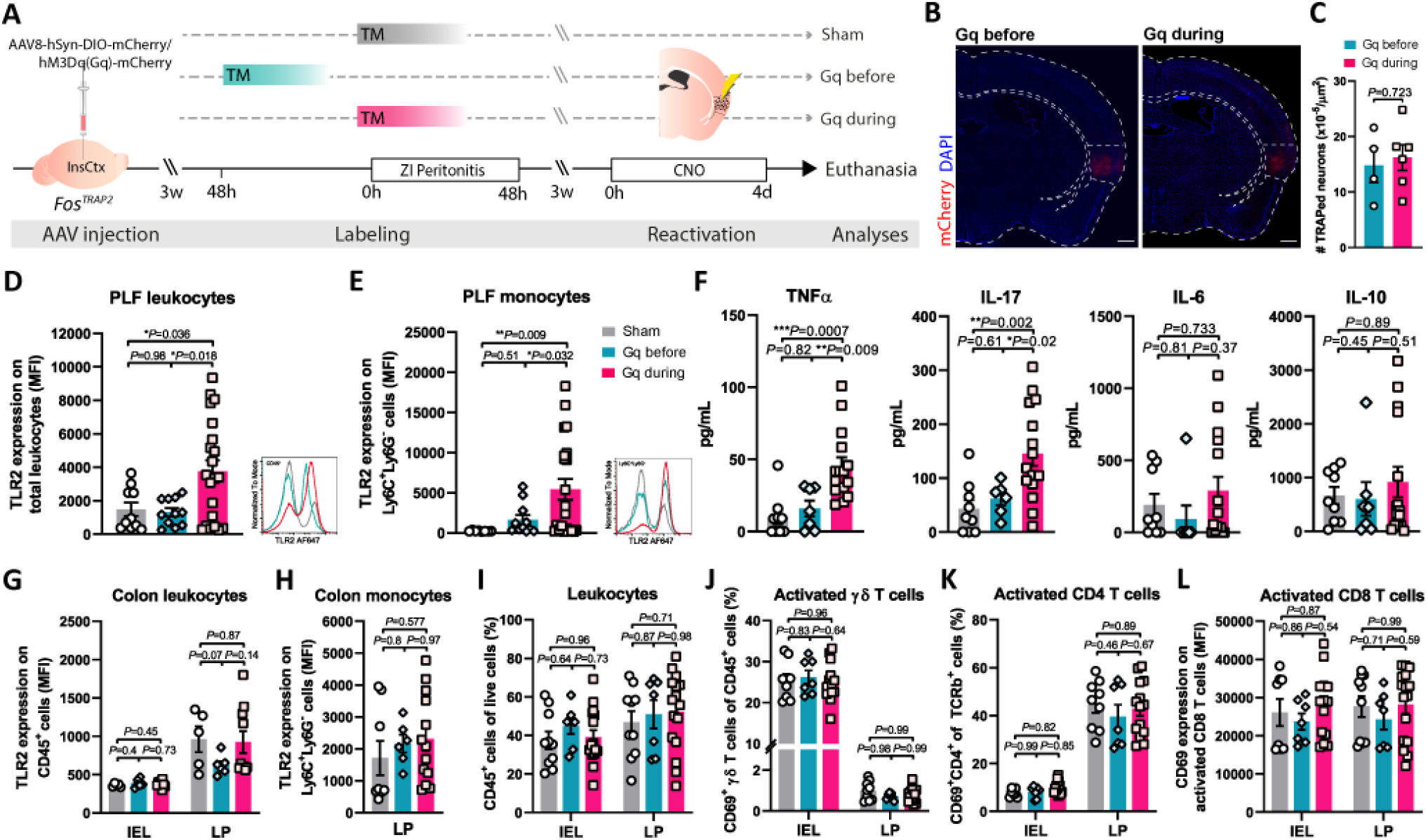
Reactivation of neuronal ensembles captured during peritonitis induces immune activation in the peritoneum but not in the colon. (**A**) Schematic representation of enabling DREADD expression in neuronal ensembles that were active before, during or after ZI-peritonitis in the InsCtx of *Fos^TRAP2^* mice (via different timing of TM injections, relative to Zymosan administration). A sham viral vector control group, expressing only the fluorescent reporter in a Cre-dependent manner, was injected with TM during ZI-peritonitis development. Three weeks after Zymosan administration, all mice received a single i.p. CNO injection, and the immune response was analyzed 24h later. (**B**) Representative fluorescence microscopy imaging showing the expression of the injected Cre-dependent DREADD (mCherry, in red) in the InsCtx in both Gq-expressing groups. Scale bar: 250 μm. (**C**) Quantification of DREADD-expressing neurons in the InsCtx of both groups. Data are represented as mean±s.e.m from *n*= 4, 6 mice; Student’s *t*-tests. (**D-E**) Graphs derived from flow cytometry analysis, showing TLR2 expression on peritoneal (**D**) leukocytes (CD45^+^ cells; *n*=10, 11, 21), and (**E**) monocytes (CD45^+^Ly6C^+^Ly6G^-^ cells; *n*=8, 11, 21) in both Gq groups, labeled before (Gq before) and during (Gq during) inflammation, and their sham viral vector controls (Sham). Data are represented as mean±s.e.m and representative histograms of TLR2 expression are presented for each graph; One-way ANOVA and Multiple *t*-tests. Two (**D**) and four (**E**) samples were excluded from the data; ROUT method for outlier detection (Q=1%). (**F**) Graphs derived from cytokine ELISA showing concentrations of TNFα (*n=*9, 7, 15), IL-17 (*n=*9, 6, 15), IL-6 (*n=*9, 7, 15), and IL-10 (*n=*9, 7, 15) in PLF of the Gq groups compared to their sham viral vector controls. Data are represented as mean±s.e.m; One-way ANOVA and Multiple *t*-tests. One sample (IL-17) was excluded from the data; ROUT method for outlier detection (Q=1%). (**G-L**) Graphs derived from flow cytometry analysis showing TLR2 expression on (**G**) IEL and LP leukocytes (*n*=5, 6, 10) and (**H**) LP monocytes (*n*=8, 6, 13), and percentages of colonic (**I**) leukocytes (CD45^+^ cells, *n*=10, 7, 16), (**J**) activated ɣɗ T cells (*n*=8/9, 7, 14/16) (**K**) activated CD4^+^ T cells (*n*=8/9, 7, 14), and (**L**) CD69 expression (MFI) on CD8^+^ T cells (*n*=8/10, 7, 14/16) of the two Gq groups and their sham viral vector controls. Data from individual mice are shown and values are represented as mean±s.e.m; One-way (**H**) and Two-way (**G, I-L**) ANOVA and Multiple *t*-tests. ELISA, enzyme-linked immunosorbent assay; InsCtx, insular cortex; TM, Tamoxifen; CNO, clozapine-*N*-oxide; PLF, Peritoneal lavage fluid; TLR2, Toll-like receptor 2; MFI, median fluorescence intensity; IEL, intraepithelial lymphocytes; LP, lamina propria. Data represent at least two pooled independent repeats.

In this model, the immunological changes were reminiscent of the inflammatory response directly induced by i.p. Zymosan administration (**Figure S4A-C**). For example, expression of Toll-like receptor 2 (TLR2) on all peritoneal leukocytes increased significantly in the Gq-during group, compared to the Gq-before and sham groups (*P*=0.0074; **Figures 5D, S4C, S5**), mainly on peritoneal monocytes (*P*=0.0137; **Figure 5E, S4C, S5**). This effect is especially relevant since TLR2 recognizes Zymosan, and is overexpressed on monocytes following i.p. Zymosan administration ^46,47^. Moreover, we found a significant increase in the levels of the cytokines TNFα (*P*=0.0004) and IL-17 (*P*=0.0035) in the peritoneal fluid of the Gq-during group compared to both the Gq-before and sham-viral vector control groups, while no significant changes were observed in IL-6 (*P*=0.385) and IL-10 (*P*=0.687) concentration (**Figure 5F**). Importantly, these effects were distinct from the effects detected in the DSS model (**Figure S3C**). Moreover, as in the case for DSS-induced colitis, general activation of the InsCtx, using TRAP-independent Gq expression, did not induce the effects observed in the peritoneum (**Figure S6A-D**), further supporting the specificity of the information encoded by defined neuronal representations.

Finally, to determine whether the immune-related information encoded in the brain also included anatomical information, we evaluated the immune response in additional tissues. We found that reactivation of neuronal ensembles captured during ZIP were specific to the peritoneum. We did not observe any changes in immune parameters in the colon (**Figure 5G-L**), mesenteric lymph nodes (mLN), or blood **(Figure S4C)**. These results demonstrate that even without an additional immune challenge, the immune-related information encoded in the InsCtx is sufficient to induce anatomically defined, specific peripheral immune reactions.

Taken together, these findings offer a new perspective on the concept of memory in the context of the immune system. To the best of our knowledge, this is the first demonstration of neuronal ensembles in the brain that acquire and retrieve specific immune-related information. In the field of neuroscience, this type of information encoding is reminiscent of memory traces ^48,49^. In recent years, such memory traces were demonstrated by retrieval of fear-related behavior with the reactivation of hippocampal neuronal ensembles captured during fear conditioning ^50–52^. Similarly, in our paradigm, reactivation of neurons captured under a unique inflammatory condition was sufficient to induce a related inflammatory response.

However, in contrast to the classical neuroscience concepts, in the context of an immune representation in the brain, there are additional potential interpretations of our data that require further investigation and will likely reveal additional mechanistic aspects of this brain-body connection. For example, it is possible that the information that was encoded by the neuronal ensembles reflects sensory input from the tissue (gut or peritoneum) rather than a memory trace of the relevant immune response. Namely, the brain did not encode a specific immune reaction, but rather a representation of the damage caused to the tissue. In this case, the reactivation of the neuronal ensembles replays the sensory inputs from the tissue, inducing a specific immune response. Such a phenomenon is in itself novel, yet distinct from the classical concept of memory. This question may be addressed, in part, by further characterizing the neuronal populations that respond to the immune activity, and the changes that they undergo (both at the cellular and network level). Moreover, elucidating the mechanism that enables the brain to capture and reactivate specific immune reactions requires an understanding of which cells or molecules in the periphery can report to the brain. Although such bidirectional communication was suggested before ^11,53,54^, it is also not clear to what extent the nervous system can communicate directly with immune cells in the periphery. It is possible that the signal from the nervous system is mediated via the tissue’s parenchymal cells (e.g., colon epithelia), which may also contribute to the anatomical specificity of the induced immune response.

We do not know how the information from the brain is transmitted to the different peripheral organs. The InsCtx is connected to all major hubs of communication with the periphery, namely the endocrine, sympathetic and parasympathetic pathways ^21,36^. Although the spatial specificity of the observed effects supports a neuronal route of communication, the underlying mechanisms must still be elucidated. Moreover, the InsCtx is known to be involved in pain processing. Does the interplay between pain and immune reactions contribute the phenomenon observed in these experiments?

In this study, we focused on the InsCtx, but this brain region is most likely only a part of the neuronal representation of the immune-related information in the brain. Other brain regions (e.g., hippocampus, amygdala) potentially encode other aspects of the immune response. In fact, it is possible that the differences between the original inflammatory responses (DSS-induced colitis and ZIP) and the response induced by neuronal reactivation (**Figure S3C and S5C**), may be attributed to the fact that we have captured only a fraction of the representation of these responses in the brain, as demonstrated in Figure 1C. Moreover, some of these observed differences may reflect the relative contribution of neuronal inputs compared to innate regulatory signaling of the immune system.

An even more fundamental question is the nature of the evolutionary advantage for the organism in encoding such detailed and specific immune information. One possibility is that the organism, which constantly records external cues (e.g., place, odor) also records its own response to these experiences as a means to enable a more effective reaction to recurring stimuli. Nevertheless, such a potentially beneficial physiological response could also lead to maladaptive conditions. For example, it was shown almost 150 years ago that presenting patients allergic to pollen with an artificial flower, is sufficient to induce an allergic response ^55^. Moreover, many gut-related disorders are suggested to be psychosomatic in etiology, induced by emotionally-salient experiences, with limited understanding of the underlying etiology ^56–58^. Our study adds a new perspective to the understanding of these pathological conditions, and presumably, a new avenue for therapeutic interventions.

## Acknowledgements

We would like to thank J. Gross, R. Yifa, Y. Berger and O. Barak for helpful discussions and technical support. We would like to thank S. Schwarzbaum for editing the manuscript and her insightful comments. We are grateful to A. Grau, O. Shenker, E. Suss-Toby, and Y. Sakoury from the Biomedical Core Facility at the Technion Faculty of Medicine for technical support. A.R is a Howard Hughes Medical Institute HHMI-Wellcome trust international scholar.

## Author Contributions

T.K designed and carried out all the experiments, interpreted the results, and wrote the manuscript. M.K, N.B, H.D and H.H contributed to the experimental design, execution of the experiments and contributed to data analysis. T.B.S, contributed to the development of the experimental tools and original experimental design. E.A, M.A, M.S, B.K, H.H and D.F contributed to the execution of the experiments. F.H and K.R contributed to the experimental design and the interpretation of the results. A.R contributed to the experimental design, interpretation of results, and wrote the manuscript. All authors commented on the final manuscript.

## Data availability

All data are provided in the main text or the supplementary materials.

## Supplementary Materials

Materials and Methods

Figures S1-S6

