## Supplementary materials for "Remembering immunity: Neuronal ensembles in the insular cortex encode and retrieve specific immune responses"

#### Materials and Methods

**Mice**. Male adult (8-12 weeks of age; 20-25 g) $Fos^{TRAP1}$ (Jackson Laboratory; B6.129(Cg)-$\textit{Fos}^{tm1.1(cre/ERT2)Luo}$/J; stock #021882), $Fos^{TRAP2}$ (Jackson Laboratory; $\textit{Fos}^{tm2.1(icre/ERT2)Luo}$/J; stock #030323) and Ai14D (Jackson Laboratory; B6;129S6-Gt(ROSA)26$Sor^{tm14(CAG\text{-}tdTomato)Hze}$/J; Stock #007908) transgenic mice were used for the activity-dependent labeling experiments. C57BL/6J mice (Jackson Laboratory; C57BL/6J; stock #000664) were used for the characterization of the DSS-induced colitis, Zymosan-induced peritonitis and the Cre-independent InsCtx activation experiments. Mice were maintained under specific-pathogen-free (SPF) conditions on a 12 h light: 12 h dark cycle (lights on at 07:00) with food and water ad libitum. All experiments were performed in accordance with the National Institutes of Health Guide for the Care and Use of Laboratory Animals [59]. All procedures and protocols were approved by the Technion Administrative Panel of Laboratory Animal Care. Throughout the experiments, mice were randomly assigned to experimental groups.

**Stereotactic injections**. Mice were anesthetized with a ketamine-xylazine mixture (ketamine 80 mg per kg body weight (mg/kg); xylazine 15-20 mg/kg; Sigma-Aldrich) in sterile saline (0.9% NaCl). After skull exposure, mice were fixed in the stereotactic frame (Stoelting) and carefully drilled through the skull, just above the target injection site. The needle was slowly lowered into the brain and left in place for 5 minutes, both before and after the injection. An adeno-associated virus 8 (AAV8)-based construct (ELSC Vector Core Facility) was used to induce Cre-dependent DREADD expression (AAV8-hSyn-DIO-hM3D(Gq)-mCherry) in active neurons, by injection of 0.40 μl (0.07 μl per min) into the right posterior InsCtx (AP,

−0.35 mm; ML, 4.0 mm; DV, 3.83 mm; relative to bregma). Control mice were injected with a sham AAV8-hSyn-DIO-mCherry construct, lacking the DREADD gene. Mice showing signs of physical distress and pain were excluded from the experiment. Experiments were performed at least 21 days after surgery to ensure the expression of DREADDs and to avoid analyzing the immediate immune response that could be induced by the virus injection. Stereotactic injection sites were verified using fluorescence microscopy on brain tissue cryosections.

**DSS-induced colitis.** For the induction of colon inflammation (colitis), mice were administered 3% DSS (dextran sulfate sodium, TdB consultancy) in their drinking water for 84 hours. This duration of DSS administration was chosen after calibration experiments and previous studies [60,61], indicating that this time point was long enough for mice to exhibit signs of colon inflammation and yet short enough to allow complete recovery within 4 weeks. Fresh DSS solutions were prepared every 2 days. During the administration period, mice were monitored for their general condition: body weight, stool consistency, and signs of illness including hunched posture and dehydrated eyes and skin. Since the acute colitis was induced only for 84 hours, there were no mice showing any severe physical distress throughout the experiments.

**Zymosan-induced peritonitis.** For peritonitis induction in experimental mice, we used Zymosan, prepared from *Saccharomyces cerevisiae* cell wall (Sigma-Aldrich). Twenty mg of Zymosan were dissolved in 1 ml of sterile HBSS (without $Ca^{+2}$ and $Mg^{+2}$, Biological Industries), during heating in a boiling water bath for 20 min. We sonicated the solution for 20 min, washed twice with HBSS (1200 rpm, 10 min) and resuspended in HBSS to a concentration of 20 mg/ml. The Zymosan solution was aliquoted and kept at -80°C. Upon use, the Zymosan solution was diluted in sterile PBS to a concentration of 20 μg/ml and a volume of 0.5 ml was administered to each mouse via intraperitoneal (i.p.) injections (10 μg of Zymosan per peritoneal cavity). As previously described [62], this dose is sufficient to induce a well-characterized inflammatory response, which is confined to the peritoneal cavity, and resolves within 48 hours after induction.

**Activity-dependent cell labeling.** Tamoxifen (Sigma-Aldrich) was dissolved in corn oil (Sigma-Aldrich) to a concentration of 20 mg/ml and kept at 4°C until use (up to 12 hours). To enable a Cre-dependent expression of the target gene (Gq-DREADD or fluorescent reporter) in active neurons, 15 mg/kg of the Tamoxifen solution was injected i.p., 24 hours prior to the immunological event of interest (i.e., colitis or the peak of peritonitis development).

**Chemogenetic neuronal activation.** To reactivate the captured neurons in the InsCtx, all mice were i.p. injected with CNO (1 mg/kg; Sigma-Aldrich) in sterile saline, 4 weeks after Tamoxifen administration. In the DSS experiments, CNO was injected for 4 days at 24-hour intervals; in the Zymosan experiments, a single dose of CNO was administered 24 hours prior to euthanizing the mice. Of note, in all experiments, mice in all groups were injected with CNO to control for the potential effects of CNO, which in mice does not convert to clozapine [63].

**Tissue preparation and immunofluorescence**. Validation of the virus injection site, evaluation of DREADD expression, analysis of neuronal phenotype in the brain and validation of flow cytometry results in colonic tissues were performed by immunofluorescence staining and fluorescence microscopy. Mice were euthanized, and their brains and 1cm of their distal colon were extracted and fixed in 4% paraformaldehyde (PFA) for 48 h, and cryoprotected in 30% sucrose solution (72 h). Coronal cryosections were sliced at 12 µm thickness and mounted on super-frost slides (Thermo scientific). For immunofluorescence, the sliced tissue sections were incubated in a blocking buffer (0.2% Triton X-100, 0.05% sodium azide, 4% bovine serum albumin in PBS) for 2 hours at room temperature. Primary antibodies diluted in blocking buffer were added to the desired concentration and the slides were incubated at 4°C overnight. The tissue sections of the brain were stained using mouse anti-NeuN (1:400, Sigma-Aldrich, MAB377), rabbit anti-GFAP (1:250, Agilent, ZO334) and rat anti-CD11b (1:250, BioLegend, 101228). The tissue sections of the colon were stained with rat anti-CD45 (1:250, BioLegend, 103128). The slides were then washed in PBS 3 times for 15 min, and a fluorescent-conjugated secondary antibody: Alexa Fluor 488-donkey anti-mouse IgG (1:500, Jackson ImmunoResearch laboratories, 715-545-151); Cy5-goat anti-rabbit IgG (1:300, Jackson ImmunoResearch laboratories, 111-175-144); Cy5-goat anti-rat

IgG (1:300, Jackson ImmunoResearch laboratories, 112-175-167 ); or Alexa Fluor 568-goat anti-rat IgG (1:300, Invitrogen, A11077), was applied for 60 min at room temperature. Slides were then washed in PBS, mounted with DAPI-mounting medium (SouthernBiotech) and covered with a glass cover slip (Bar-Naor). To evaluate c-Fos expression, mice were sacrificed 90 min after CNO injection and treated as described above. Fixed sections were stained with rabbit anti-c-Fos antibody (1:2000, abcam, ab190289). We quantified the number of the c-Fos$^{+}$ cell nuclei, normalized to the total number of mCherry-expressing cells. All images were taken at 10× or 20× magnification using an Axio imager M2 microscope (Carl Zeiss Inc. US). The quantification of positive or double-positive cells was performed using ImageJ (fiji) Software.

**Flow cytometry.** Mice were euthanized, and their blood, mesenteric lymph nodes (mLN), peritoneal lavage fluid (PLF) and colon (starting at the cecocolic junction proximally and ending at the anus distally) were collected. EDTA-coated tubes were used for blood collection, and centrifuged at 1200 rpm for 10 min. For blood cell analysis, 1 ml of blood was incubated with 9 ml of RBC Lysis buffer (BD biosciences, 555-899) for 15 min and washed twice with PBS. mLN were dissociated in PBS into single-cell suspensions and mesh-filtered (40μm) to remove fat tissue and debris. Peritoneal cells were collected via peritoneal lavage; 6 ml of cold PBS was carefully injected into the peritoneum and the cell-containing fluid was collected with a Pasteur pipette into 15 ml tubes. These were centrifuged at 1200 rpm for 5 min. PF supernatant was extracted and frozen at -20$^{o}$C for further protein analysis. The recovered cells were incubated for 3 minutes with 1 ml of Lysis buffer and washed once with PBS. In case of blood-contamination in PF samples, these samples were excluded from the experiment. Colon tissue was mechanically dissociated into intraepithelial lymphocytes (IEL) and enzymatically dissociated into lamina propria (LP) single-cell suspensions using the Lamina propria dissociation kit (Miltenyi Biotec, 130-097-410). For viability staining, cells ($10^{6}$) were washed once with PBS, resuspended in Zombie Aqua$^{TM}$ dye solution (1:000, Biolegend, 423106) and incubated in the dark for 15 min at room temperature. For extracellular staining, cells were washed with FACS staining buffer (PBS containing 1% bovine serum albumin, 1 mM EDTA and 0.05% sodium azide) and incubated with antibodies for 30 min at 4 °C. For intracellular staining, the samples were first stained for extracellular markers as described above, fixed and permeabilized with BD Cytofix/Cytoperm kit, and stained with the intracellular antibodies. The following mAbs were used: Alexa-fluor-700 anti-CD45 (BioLegend, 103128), Brilliant-violet-510 anti-Ly-6C (BioLegend, 128033), Alexa-fluor-647 anti-TLR2 (BioLegend, 121809), PerCP anti-CD8a (BioLegend, 100732), Biotin anti-CD69 (BioLegend, 104504) together with Alexa-fluor-488-conjugated streptavidin (Jackson, 016-540-084), PE/Cy7 anti-IFN-γ (Biolegend, 505826), PerCP/Cy5.5 anti-CD11b (BioLegend, 101228), APC anti-F4/80 (BioLegend, 123116), FITC anti-Ly6G (BioLegend, 127606), APC anti-CD4 (BioLegend, 100412), PE/Cy7 anti-CD11c (BioLegend, 117318), Alexa-fluor-488 anti-I-A/I-E (BioLegend, 107616), PE anti-TCRγδ (Biolegend, 118108), Pacific-blue anti-TCRβ (BioLegend, 109226), Brilliant-violet-605 anti-CD103 (BioLegend, 121433). All antibodies were validated by the manufacturers for flow application, as indicated on the manufacturers' websites. The samples were re-suspended in 250 μl of 1% PFA and analyzed by flow cytometry. Samples were analyzed with a CytoFLEX S cell analyzer and FlowJo software.

**Measurement of colon and peritoneal lavage fluid cytokine levels.** For peritoneal lavage fluid (PLF) protein analysis, 6 ml of cold PBS was carefully injected into the peritoneum and the cell-containing fluid was collected with a Pasteur pipette into 15 ml tubes. These were centrifuged at 1200 rpm for 5 min. PLF supernatant was extracted and frozen at -20$^{o}$C for further protein analysis. For colon protein analysis, a 200 mg piece of the tissue was collected from each mouse. These samples were homogenized in a homogenization buffer (1:2, mg/μl) using stainless steel beads and the Bullet Blender Storm 24 (Next Advance), according to the recommended protocol at the manufacturer's website. PF supernatant and homogenized colon samples were analyzed using standard ELISA kits for TNF-α, IL-6, IL-17 and IL-10 (PeproTech).

**Statistical analysis.** In all experiments, significance levels of the data were determined using Prism5 (GraphPad Software). Experiments were analyzed by two-tailed unpaired or paired Student’s *t*-test or One-way ANOVA and multiple *t*-test, with a significance threshold set at $P$=0.05. Data are represented as mean

± standard error of mean (s.e.m). Outliers were excluded using ROUT method for outlier identification [64] with a False Discovery Rate less than 1% (Q<1%). All experiments were performed at least twice.

### Supplementary figures

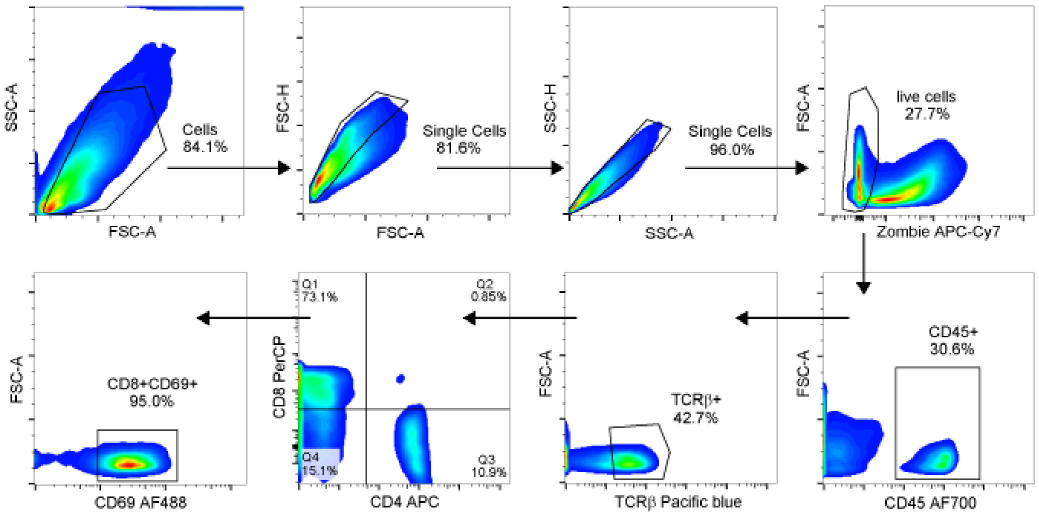


**Figure S1. Immune cells in the colon following reactivation of InsCtx neuronal ensembles captured during colitis.** Gating strategy of the different immune cell subpopulations in the colon mucosal layer analyzed by flow cytometry and presented in Figure 2D-G.

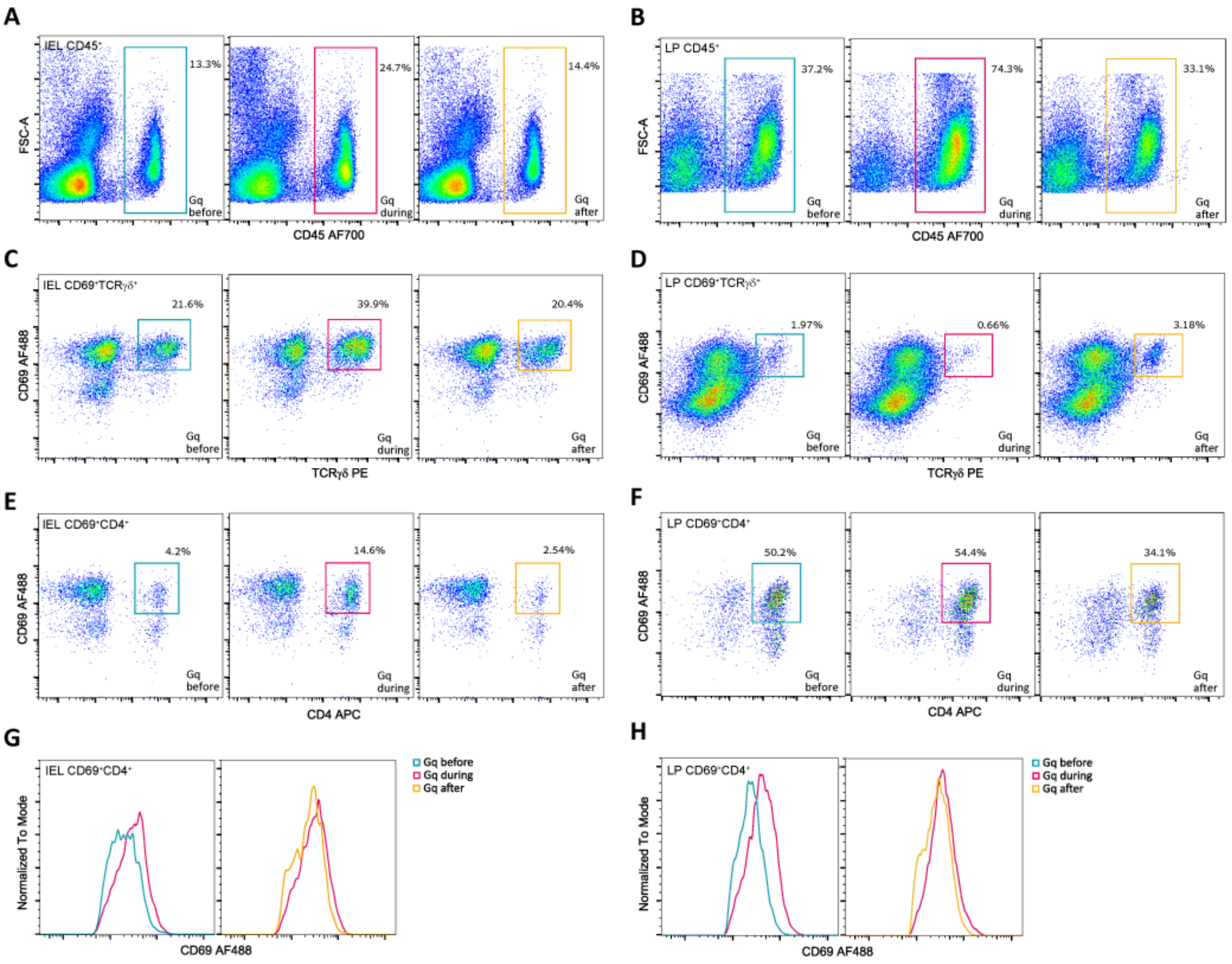


**Figure S2. Immune cells in the colon following reactivation of InsCtx neuronal ensembles captured at different time-points relative to colitis induction.** (**A-F**) Representative dot plots of mucosal (**A, B**) leukocytes ($CD45^+$ cells), (**C, D**) activated γδ T cells and (**E, F**) activated CD4 T cells from IEL and LP of the Gq-before, during and after groups (gated in blue, red and orange, respectively). (**G, H**) Representative histograms of CD69 expression level (MFI) on activated CD4 T cells from (**G**) IEL and (**H**) LP of Gq-before (blue), Gq-after (orange) groups layered against histograms of the Gq-during group (red). Experimental groups are depicted in Fig. 3A. InsCtx, insular cortex; IEL, intraepithelial lymphocytes; LP, lamina propria; MFI, median fluorescence intensity.

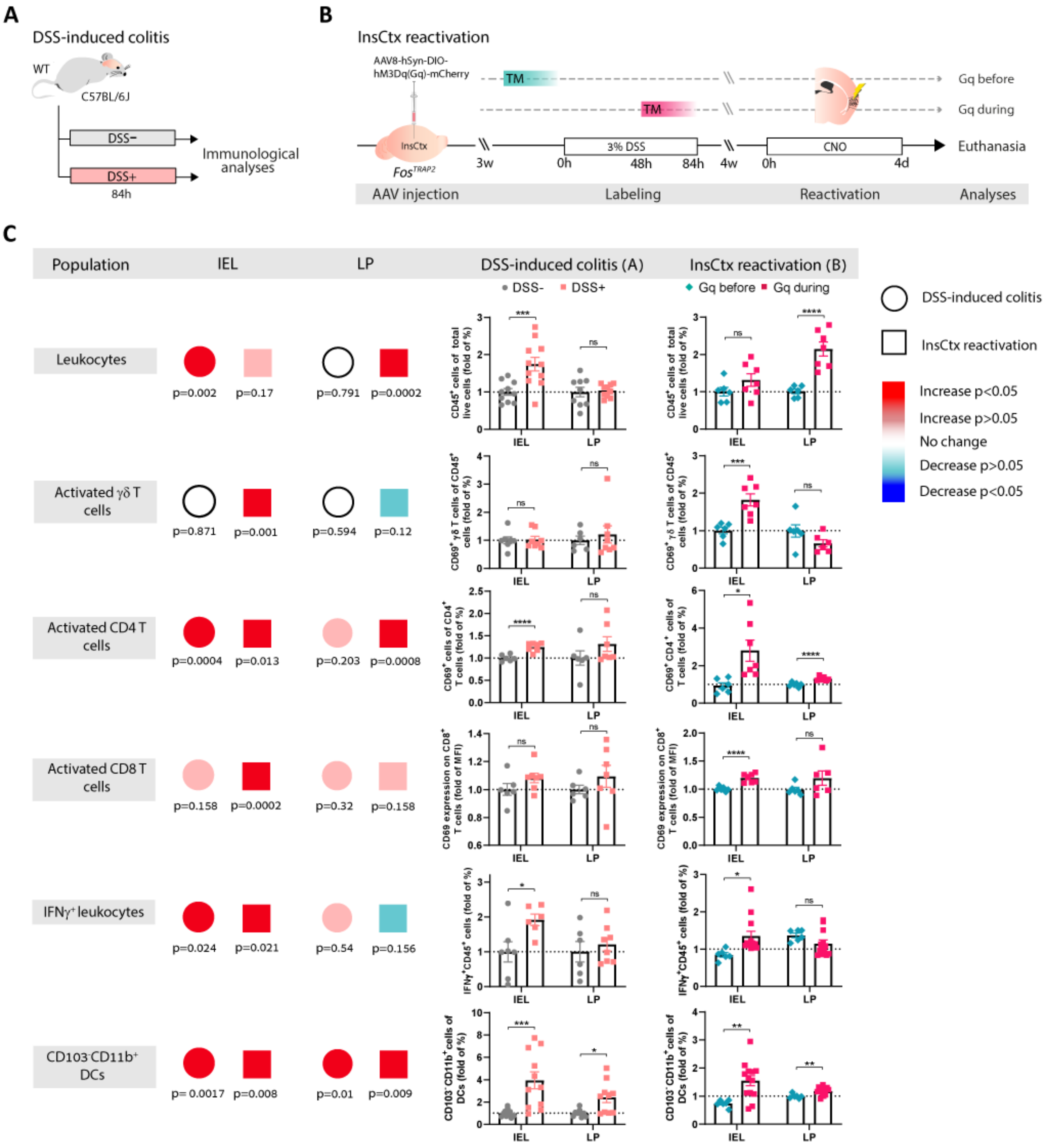


**Figure S3. Comparison between immune characteristics induced by DSS administration alone and reactivation of InsCtx neuronal ensembles captured during colitis.** (**A**) Schematic representation of the acute DSS-induced colitis model in C57BL/6J WT mice. Mice were administered with 3% DSS in their drinking water (DSS+) for 84h (simulating the time-period, in which *Fos*$^{TRAP2}$ mice received DSS during neuronal capturing). As controls, a second group of mice was administered with normal drinking water

(DSS-). (**B**) Schematic representation of capturing neuronal ensembles before (Gq-before) or during (Gq-during) DSS-induced colitis in *Fos*$^{TRAP2}$ mice. These neuronal ensembles in the InsCtx were then reactivated and the immune response was analyzed. TM indicates the timing of tamoxifen administration. (**C**) Summary table of the differences and similarities in the cellular immune response in the mucosal layer of the colon between the two experiments (DSS-induced colitis vs. InsCtx reactivation). Presented are the immune cell populations (Population) that have changed between the two experimental groups in each experimental design (DSS+ relative to DSS- and Gq-before relative to Gq-during) in either of the colonic mucosal layers (IEL or LP). Large circles indicate the DSS-induced colitis and the squares indicate the InsCtx neuronal activation paradigm. Their colors indicate the trend of change (increase or decrease) and its significance level is indicated below ($P<0.05$ or $P>0.05$). Graphs presented on the right panel are derived from flow cytometry analysis showing fold change of leukocytes ($CD45^+$ cells), activated γδ T cells and activated $CD4^+$ T cells percentages, $CD8^+$ T cells extent of activation, and $IFN\gamma^+$ leukocytes and $CD103^-$ $CD11b^+$ dendritic cells (DCs) percentages. Data from individual mice are shown and values are represented as mean±s.e.m; ns, not significant, ***$P<0.005$, ****$P<0.0005$; Multiple *t*-tests. InsCtx, insular cortex; WT, wild-type; DSS, dextran sulfate sodium; TM, Tamoxifen; CNO, clozapine-*N*-oxide; IEL, intraepithelial lymphocytes; LP, lamina propria; MFI, median fluorescence intensity; DCs, dendritic cells. Data represent two pooled independent repeats for each experimental design.

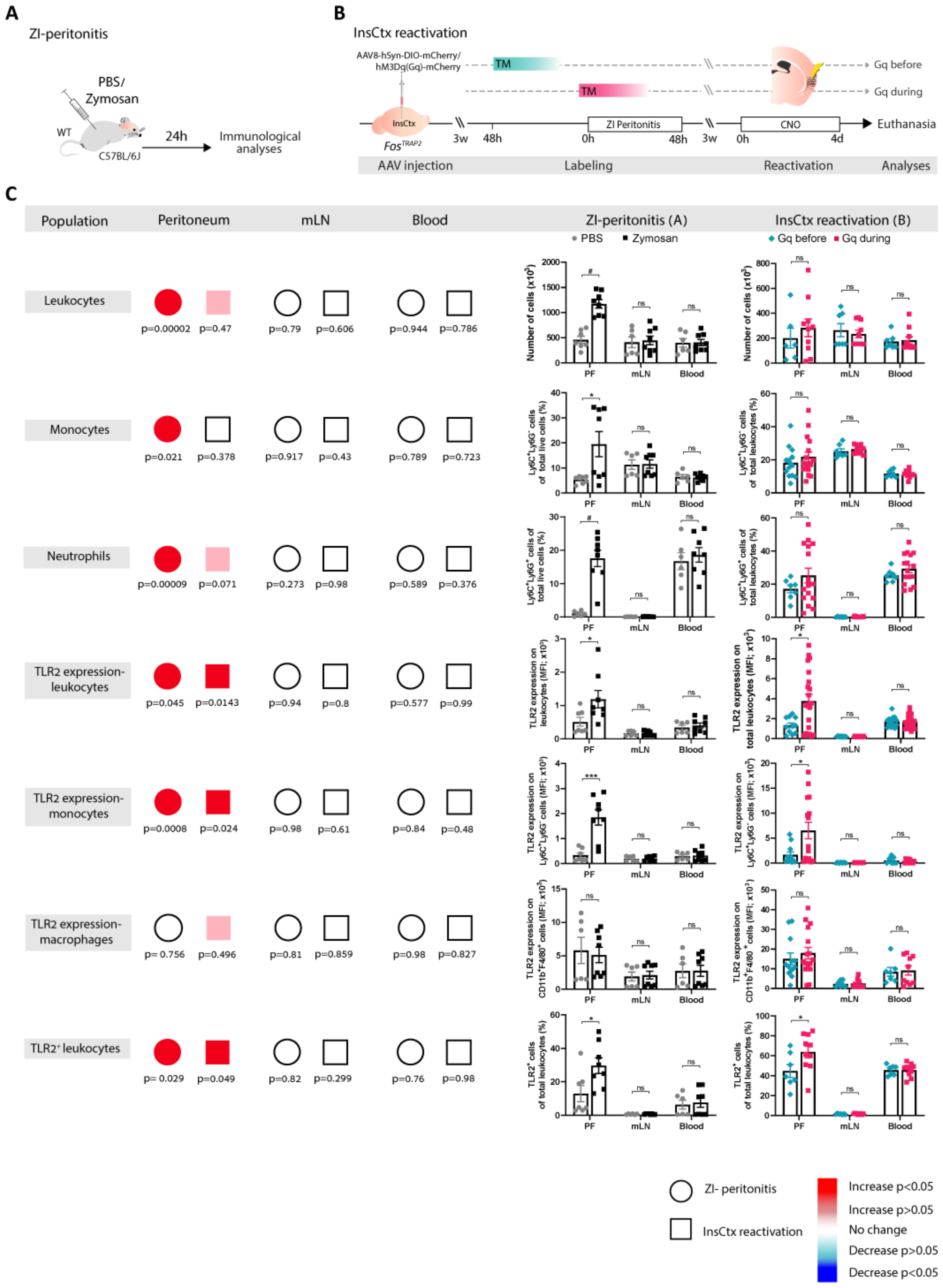
A
ZI-peritonitis
PBS/
Zymosan
WT
C57BL/6J
24h
Immunological analyses
B
InsCtx reactivation
AAV8-hSyn-DIO-mCherry/
hM3Dq(Gq)-mCherry
TM
TM
Gq before
Gq during
InsCtx
Fos^TRAP2
3w
48h
ZI Peritonitis
0h
48h
3w
CNO
0h
4d
Euthanasia
AAV injection
Labeling
Reactivation
Analyses
C
Population
Peritoneum
mLN
Blood
ZI-peritonitis (A)
InsCtx reactivation (B)
PBS
Zymosan
Gq before
Gq during
Leukocytes
p=0.00002 p=0.47 p=0.79 p=0.606 p=0.944 p=0.786
Monocytes
p=0.021 p=0.378 p=0.917 p=0.43 p=0.789 p=0.723
Neutrophils
p=0.00009 p=0.071 p=0.273 p=0.98 p=0.589 p=0.376
TLR2 expression-leukocytes
p=0.045 p=0.0143 p=0.94 p=0.8 p=0.577 p=0.99
TLR2 expression-monocytes
p=0.0008 p=0.024 p=0.98 p=0.61 p=0.84 p=0.48
TLR2 expression-macrophages
p= 0.756 p=0.496 p=0.81 p=0.859 p=0.98 p=0.827
TLR2⁺ leukocytes
p= 0.029 p=0.049 p=0.82 p=0.299 p=0.76 p=0.98
PF
mLN
Blood
ZI- peritonitis
InsCtx reactivation
Increase p<0.05
Increase p>0.05
No change
Decrease p>0.05
Decrease p<0.05

**Figure S4. Similar immune characteristics induced by Zymosan administration and reactivation of InsCtx neuronal ensembles captured during peritonitis.** (**A**) Schematic representation of the ZI-peritonitis model in C57BL/6J WT mice. Mice received a single i.p. injection of Zymosan (identical to the Zymosan injection *Fos*$^{TRAP2}$ mice received during neuronal capturing). As controls, a second group of mice was administered with a single dose of PBS i.p. injection. Twenty-four hours after injection, peritoneal lavage fluid, mLN and blood were extracted from both groups of mice and immunologically analyzed. (**B**) Schematic representation of capturing neuronal ensembles before (Gq-before) or during (Gq-during) ZI-peritonitis in *Fos*$^{TRAP2}$ mice. These neuronal ensembles in the InsCtx were then reactivated and the immune response was analyzed. TM indicates the timing of tamoxifen administration. (**C**) Summary table of the differences and similarities in the cellular immune response in the peritoneum, mLN and blood between the two experiments (ZI-peritonitis vs. InsCtx reactivation). Presented are the immune cell populations (Population) that have changed between the two experimental groups (Zymosan relative to PBS and Gq-before relative to Gq-during) in each of the experimental designs. Large circles and squares indicate the type of experiment and their colors indicate the trend of change (increase or decrease) and its significance level ($P$<0.05 or $P$>0.05). Graphs presented are derived from flow cytometry analysis, showing the number of leukocytes ($CD45^+$ cells), percentages of monocytes and neutrophils, TLR2 expression on leukocytes, monocytes and macrophages and percentages of $TLR2^+$ leukocytes. Data from individual mice are shown and values are represented as mean±SEM; ns, not significant, *$P$<0.05, ***$P$<0.005, #$P$<0.00005; Student's *t*-test. InsCtx, insular cortex; WT, wild-type; i.p., intraperitoneal; ZI-peritonitis, Zymosan-induced peritonitis; TM, Tamoxifen; CNO, clozapine-*N*-oxide; TLR2, toll-like receptor 2; PF, peritoneal fluid; mLN, mesenteric lymph-nodes; MFI, median fluorescence intensity. Data represent two pooled independent repeats for each experimental design.

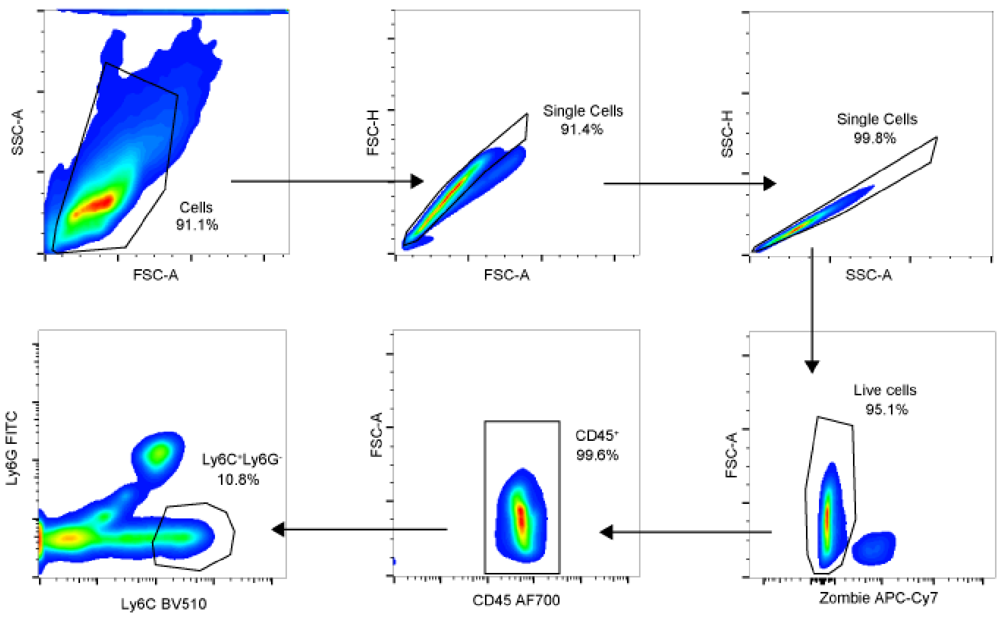


**Figure S5. Immune cells in the peritoneal cavity following reactivation of InsCtx neuronal ensembles captured during peritonitis.** Gating strategy of the immune cell subpopulations analyzed by flow cytometry and presented in Figure 5D-E.

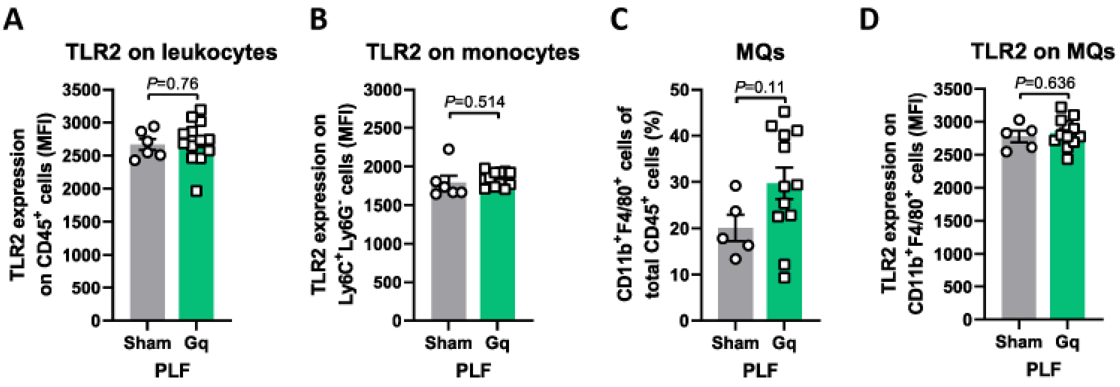


**Figure S6. Nonspecific activation of the InsCtx elicits no apparent cellular immune response in the peritoneum.** (**A-D**) Graphs derived from flow cytometry analysis showing TLR2 expression on (**A**) PLF leukocytes (*n*=6, 13) and (**B**) monocytes (*n*=6, 12), (**C**) percentages of PLF MQs ($CD45^+CD11b^+F4/80^+$ cells, *n*=5, 12), and (**D**) MQs' TLR2 expression level (*n*=5, 12) of the Gq group and its sham viral vector control. Data from individual mice are shown and values are represented as mean±s.e.m; Student's *t*-test. InsCtx, insular cortex; PLF, Peritoneal lavage fluid; TLR2, Toll-like receptor 2; MQs, macrophages; MFI, median fluorescence intensity. Data represent two pooled independent repeats.
